# A scale-invariant log-normal droplet size distribution below the critical concentration for protein phase separation

**DOI:** 10.1101/2023.04.11.536478

**Authors:** Tommaso Amico, Samuel Dada, Andrea Lazzari, Michaela Brezinova, Antonio Trovato, Michele Vendruscolo, Monika Fuxreiter, Amos Maritan

## Abstract

Many proteins have been recently shown to undergo a process of phase separation that leads to the formation of biomolecular condensates. Intriguingly, it has been observed that some of these proteins form dense droplets of sizeable dimensions already below the critical concentration, which is the concentration at which phase separation occurs. To understand this phenomenon, which is not readily compatible with classical nucleation theory, we investigated the properties of the droplet size distributions as a function of protein concentration. We found that these distributions can be described by a scale-invariant log-normal function with an average that increases progressively as the concentration approaches the critical concentration from below. The results of this scaling analysis suggest the existence of a universal behaviour independent of the sequences and structures of the proteins undergoing phase separation. While we refrain from proposing a theoretical model here, we suggest that any model of protein phase separation should predict the scaling exponents that we reported here from the fitting of experimental measurements of droplet size distributions. Furthermore, based on these observations, we show that it is possible to use the scale invariance to estimate the critical concentration for protein phase separation.

## Introduction

Many proteins have been shown to undergo a phase separation process into a liquid-like condensed state^1-5^. This process appears to be of physiological significance since it may lead to the formation of biomolecular condensates^1-7^. As a consequence, it is closely controlled by the protein homeostasis system^8,9^, and its dysregulation has been associated with a broad range of human diseases^10,11^. It is therefore important to understand the fundamental nature of this process^5,12,13^ to provide insights for the identification of ways of modulating it through pharmacological interventions.

To help understand the nature of the transition underlying protein phase separation on the basis of recent experimental observations^26,27^, here we study the distribution of the size of the droplets below the value of the concentration at which the transition occurs, which here is referred to as the critical concentration, ρ_c_. This question appears as a promising starting point to develop new insights since it has been reported that proteins can form droplets of sizeable dimensions already well below the concentration at which phase separation occurs^26,27^. This behaviour is not predicted by classical nucleation theory^28^, and not readily consistent with the idea that the protein phase separation process can be described as a first-order phase transition. This is because, in a first-order phase transition, nucleation takes place in a supersaturated system^29^, while in a subsaturated system particles can still self-assemble, but with a probability that decreases exponentially with the size of the assemblies.

In the present study, we asked whether the experimental data on the droplet size distributions obey scale invariance, a general characteristic of complex systems, which is often used to reveal universal patterns underlying self-assembly. We report the observation that the droplet size distributions of the proteins FUS and α-synuclein follow a scale-invariant log-normal behaviour. These findings are consistent with a universal behaviour resulting from the presence of an increasingly large correlation length, ξ, as the concentration approaches the critical concentration from below. The correlation length is an emergent characteristic, and it is related to the typical spatial range over which density fluctuations are correlated. When ξ is sufficiently large, one can expect scale invariance and finite-size scaling^30^ to occur within a range of lengths spanning from the molecular size to ξ. This means that physical observables cease to depend explicitly on numerous microscopic details that are peculiar to spatial scales smaller than ξ, thus leading to a universal behaviour characterized by quantities obtained by coarse graining over scales smaller than ξ.

At a first-order phase transition, ξ is finite, and if it is not large enough, the length range discussed above remains too short to observe scale invariance. However, the vicinity of the spinodal line, where nucleation disappears as the dilute phase becomes unstable, to the coexistence curve, where nucleation appears as the dilute phase becomes metastable, might cause a large increase of ξ as the first-order phase transition is approached by increasing the concentration. If it is not preceded by a first-order phase transition, the spinodal line would correspond to a second-order phase transition, resulting in infinite ξ, with scale invariance holding on all length scales larger than the molecular scale. Various alternative scenarios contributing to the emergence of significant correlations will be mentioned in the Discussion section. Nevertheless, our scaling analysis exhibits a high degree of generality, devoid of reliance on specific underlying models.

As an application of the above observations, we address the question of whether scale invariance holds for droplet size distributions near the coexistence curve. As a practical consequence, we use this observation to propose a procedure to overcome the challenge of estimating the critical concentration, ρ_c_, which enters explicitly into our scaling analysis. Such challenge arises from the fact that close to the critical concentration, the timescale required for the equilibration of a system grows together with ξ, thus exceeding the timescale amenable to experimental observation.

Our analysis of experimental data indicates that: (i) scale invariance does indeed hold near the coexistence curve, and (ii) the droplet size distribution is log-normal. Based on the properties of the scale-invariant log-normal distribution of droplet sizes, we investigate a correlation between the moments of the distribution and the distance from ρ_c_.

Finally, we note that methods to assess the critical concentration are crucial for understanding the location of proteins in their phase diagram, their proximity to the phase boundary between the native and droplet states, and how pharmacological interventions can modify their phase behaviour. To address this problem, we report how the moments of the distribution can be used as a scale-invariant gauge to estimate the critical concentration. In this way, an accurate estimate of the critical concentration is possible because it is based on measurements carried out away from the critical point, under conditions such that fluctuations are small, and hence experimental errors are smaller than in the proximity of the transition.

## Results

### Formulation of the scaling ansatz

Empirical evidence indicates that protein self-assembly into liquid-like condensates is characterized by: (i) a phase separation transition at a concentration ρ_c_, (ii) a formation of droplets of sizeable dimensions already below ρ_c_, and (iii) a droplet size distribution that, after an initial transient, does not change with the experimental observation time, although individual droplets can form, grow, shrink and dissolve. An initial analysis of droplet size distributions observed experimentally led us to ask whether the region near the transition could be described in terms of a scaling theory, as commonly done for critical phenomena (29), as summarized above. We also note that this approach is analogous to analyzing the cluster size distribution in a percolation problem^31^.

In our analysis, we called *P*_*>*_(*s*|*ρ*) the survival distribution function (SDF = 1-CDF, where CDF is the cumulative distribution function) corresponding to the probability to observe a droplet of size greater than *s*, when the concentration is *ρ*. The distance from the critical concentration *ρ*_*c*_ is measured in terms of the dimensionless variable 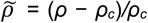, which also allows to compare data from different experiments, as explained below. *P*_*>*_(*s*|*ρ*) in general depends separately on *s, ρ* and many other parameters characterizing the process, including temperature. However, if scale invariance holds in the vicinity of *ρ*_*c*_, i.e. when 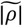 is small and the correlation length of the system is large enough, we would expect *P*_*>*_*(s*|*ρ)* to depend on *ρ* and on other details pertaining to the microscopic scales only through the characteristic droplet size, *s*_*c*_. The characteristic size *s*_*c*_ is defined, apart from a proportionality constant (see below), as the ratio of the second to the first moment of the droplet size distribution. This leads us to formulate the following scaling ansatz^31^

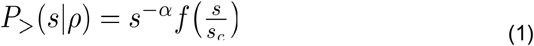

where, *s*_*c*_ depends on *ρ*, and it is expected to diverge at the critical concentration as^30,31^

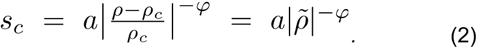

where *α* ≥ and *φ >* 0 are critical exponents, *a* is a constant and *f* is the so-called scaling function^30^.

Thus, the scaling of the SDF is equivalent to saying that, apart from the singular behaviour *s*^*− α*^, the remaining *s* and *ρ* dependence occurs only through the ratio *s/s*_*c*_. All extra dependencies are encapsulated in *s*_*c*_ through the constant *a*, ρ_c_ and, possibly, on the specific form of the scaling function *f*. A consequence of scaling and singular behaviour described by Eq. (2), i.e. the divergence of the characteristic size of the droplets, is that details of any specific system may not affect the value of the critical exponents, which are therefore expected to be universal, i.e. independent of specific details of the system.

To determine the exponents *α* and *φ*, we introduce the moments of *P*_*>*_(*s*|*ρ*) for k>0

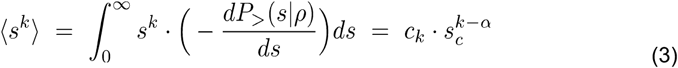

where the scaling ansatz Eq. (1) has been used in the last step and *c*_*k*_ is given by

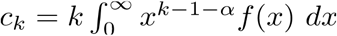

which depends on the function *f* but is independent of 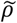. From Eq. (3), we deduce that

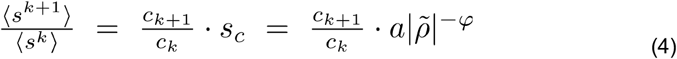

If the scaling ansatz of Eq. (1) is correct, by plotting the ratio of moments ⟨*s*^*k*+1^⟩*/*⟨*s*^*k*^⟩ for various values of *k* as a function of 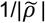 in a double logarithmic scale, we should obtain straight parallel lines with slope *φ* and intercept *ln(ac*_*k+1*_*/c*_*k*_).

### Scaling behaviour of the droplet size distributions of FUS

We investigated the validity the ansatz in Eq. (2) using experimental data on the RNA-binding protein FUS, which are available for both the untagged and the SNAP-tagged protein^27^ (**Table S1**). We calculated the survival distribution function, Eq. (2), and its moments, Eq. (3). We then plotted the moment ratios versus the inverse distance from the critical concentration 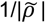 in double logarithmic scale (**Figure 1**). We used the estimates of the critical concentrations, ρ_c_ = 5.0 μM for FUS and ρ_c_ = 5.4 μM for SNAP-tagged FUS, obtained below, in a self-consistency check of the validity of the scaling ansatz. We observed that the moment ratios at different distances from the critical concentration fall onto straight lines, as predicted by the scaling ansatz. In addition, the weighted average slope (see Eqs. (15) and (16) below) of the lines for different moment ratios is 0.95 ± 0.05 for untagged FUS, and 0.95 ± 0.05 for SNAP-tagged FUS, which is in good agreement with φ=1 for the exponent in the scaling ansatz in Eq. (1) (**Figure 1A**,**C**).

**Figure 1.**
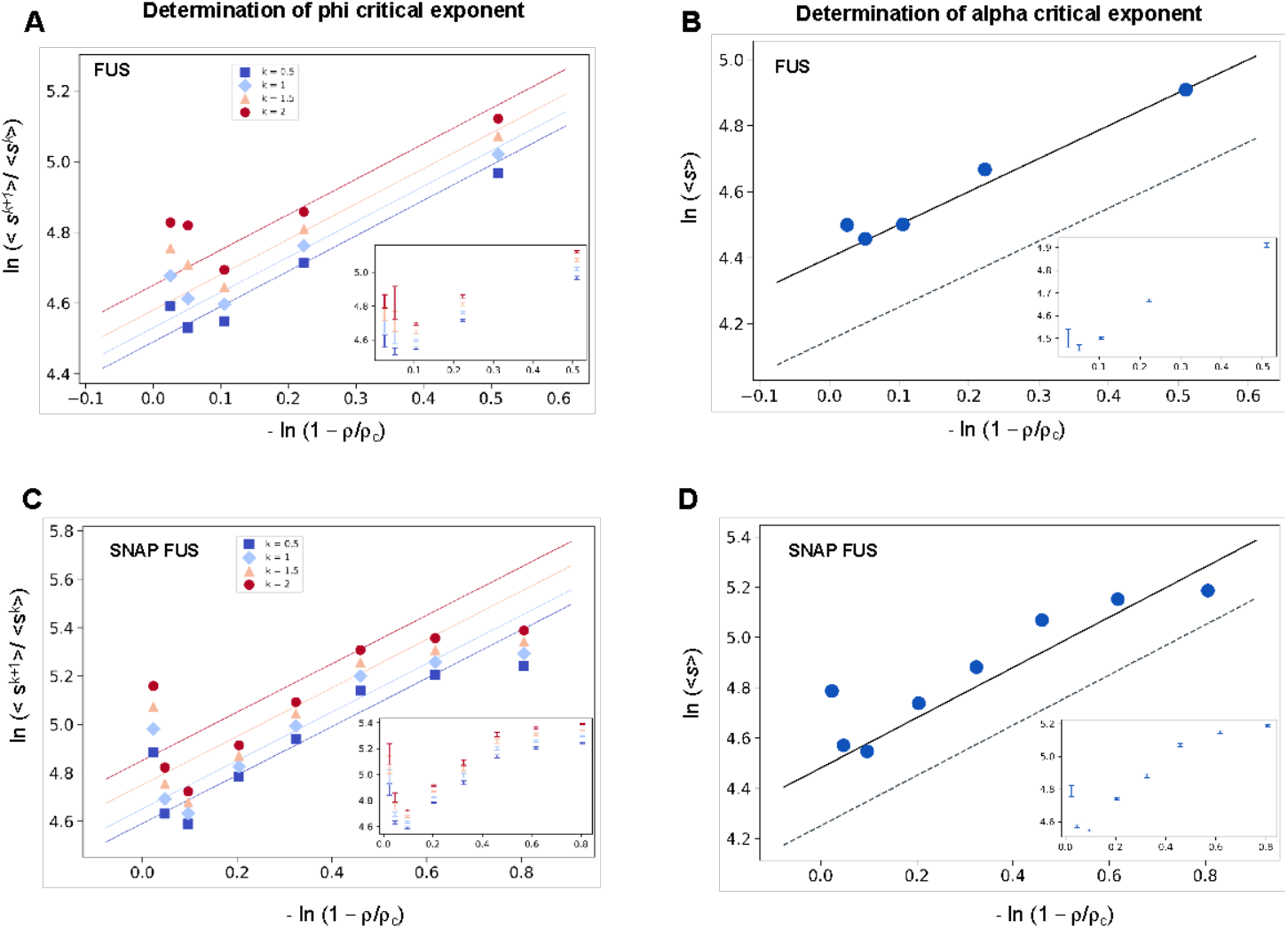
Determination of the critical exponents for FUS of the scaling invariance. **(A**,**C)** Determination of the exponent *φ* for FUS (A) and SNAP-tagged FUS (C). The ratios of the average moments of the droplet sizes (<s^k+1^>/<s^k^>, at k=0.5, 1, 1.5, 2, Eq. (4)) are represented at various distances from the critical concentration 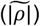. The exponent *φ* for each value of k was determined by error weighted linear regressions. The exponent *φ* and its error were determined as mean and standard deviation of the three independent measurements (Eqs. (15) and (16)). Error bars are shown in inset for graphical clarity. (**B**,**D)** Determination of the exponent *α* for FUS (B) and SNAP-tagged FUS (D). The mean of the droplet size distributions is plotted at various distances from the critical concentration 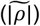. The value of the exponent *m* was determined by error weighted linear regression (Eq. (6)), using φ=1, where the errors were standard deviations of the three independent measurements (Eq. 16). Error bars, which were obtained as the standard deviation of the three independent measurements are shown in inset for graphical clarity. Error weighted linear regressions are performed in both cases excluding the data point at the lowest concentration ρ = 0.125 μM. The fit corresponding to the scaling ansatz, compatible with *φ*=1 and *α*=0, is represented by a dashed gray line with a slope of 1.

Having determined the exponent *φ*, we can also determine the exponent *α* using Eq. (3)

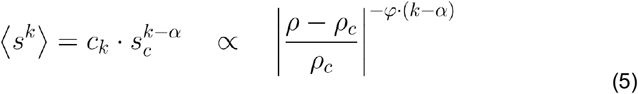

The exponent *α* is then calculated from

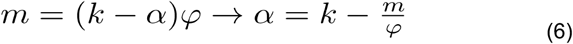

where *m* is the slope of the linear fit of the double logarithmic plot of <s> (the first moment, k=1) vs 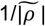. We then plotted the mean droplet size, <s>, versus the inverse distance from the critical concentration 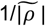 on a natural logarithm scale, which could be fitted using a line with a slope m= 0.99 ± 0.05 for untagged FUS, and 0.93 ± 0.07 for SNAP-tagged FUS (**Figure 1B,D**), which is consistent with m=1. Using the value of φ = 1, determined based on Eq. (4), we obtain α = 0. Taken together, these data support the validity of the scaling ansatz of Eq. (1).

### The droplet size distribution of FUS is log-normal

The above analysis suggests that the droplet size distribution may follow a log-normal distribution. The scaling ansatz of Eq. (1) for the SDF is equivalent to the following scaling for the probability density distribution

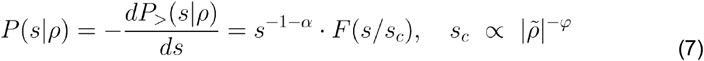

where *f*, the scaling function in Eq. (1), and *F* are related as follows

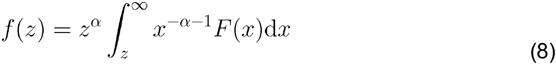

The log-normal droplet size distribution *P*(*s*|*ρ*) is

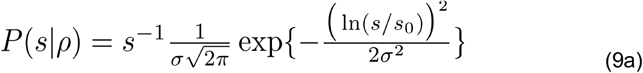

With

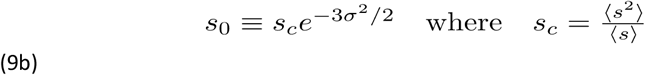

being the characteristic droplet size as defined above. Consequently, the size survival distribution function is

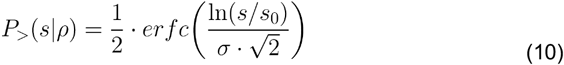

The values of *s*_*0*_ and σ can be determined as the average and the variance, of *ln(s/u)* obtained at each concentration:

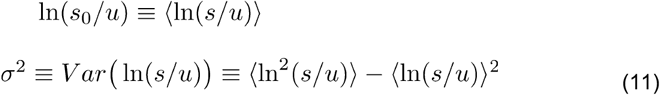

where *u* is an arbitrary (and irrelevant) constant with the same units as *s*. In the following, when not stated, it is implicitly assumed that *u=1* in the same units as *s*. The droplet sizes follow a log-normal distribution only if the survival distribution functions or, equivalently, the size distribution functions, multiplied by s, collapse when plotted versus Eq. (9) or Eq. (10) with the values of s_0_ and σ of each droplet size distributions obtained at different concentrations (Eq. (11)). We determined *s*_*0*_ and σ values for each distribution (**Figure 2A,B**) and plotted the properly rescaled size distribution functions versus [ln(s/s_0_)]/σ (**Figure 2C**). We observed that the size distribution functions collapsed for both FUS and SNAP-tagged FUS (**Figure 2C**). Furthermore, the collapsed curve overlapped with the analytic log-normal distribution we computed with s_0_=1, σ=1, the normal distribution in the rescaled variables.

**Figure 2.**
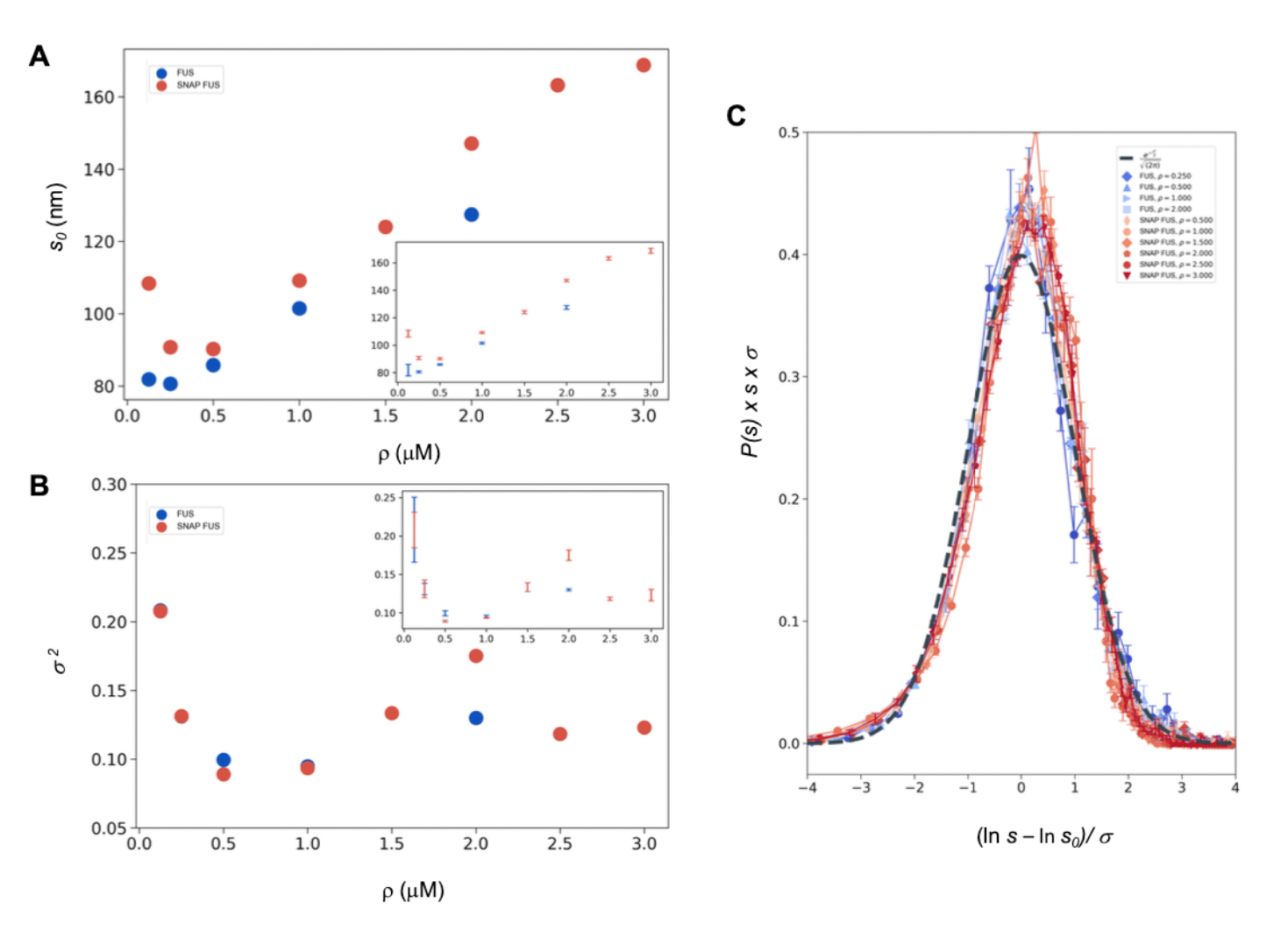
Log-normal behavior of FUS and SNAP-tagged FUS size distributions below the critical concentration. **(A, B**) Variation of the size distribution with protein concentration: ln*s*_*0*_ (A) and σ (B). ln*s*_*0*_ and σ (inset) were computed for FUS (blue) and SNAP-tagged FUS (red) using Eq. (11). Error bars (in inset for graphical clarity) are estimated from three independent measurements. While droplet sizes increase with concentration, the width of distribution does not change considerably. **(C)** The collapse of the droplet size distribution functions is consistent with a log-normal behaviour. The droplet size distribution functions for both untagged FUS (blue) and SNAP-tagged FUS (red) are plotted after rescaling the sizes by the ln*s*_*0*_ and σ values, the first and second moment of the logarithm of the droplet size distribution, which are a function of the concentration. The rescaled curves for both the untagged and the tagged protein collapse to the normal distribution (gray dashed), as expected when the non-rescaled droplet sizes follow a log-normal distribution.

The observed collapse supports the observation that droplet size distributions from different experiments follows a log-normal behaviour.

### Independence of the variance of the distribution from the concentration of FUS

The log-normal behaviour described above is consistent with the scale invariance underlying Eq. (1) if the variance of the log-normal distribution, *σ*, is independent of ρ or, equivalently, of 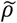 ^32^. Indeed, comparing Eq. (1) with Eqs. (7), (9a,b) and (10) we obtain that the scaling invariance holds with

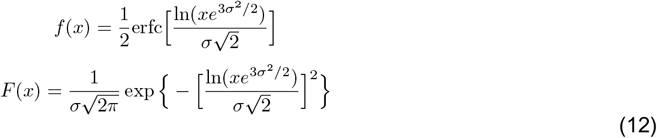

with *α=0 and φ = 1* (**Figure 1**). Notice that the log-normal distribution implies that α*=0* whereas the value of *φ* is not determined a priori. Furthermore, the *k*-th moment of the log-normal distribution is

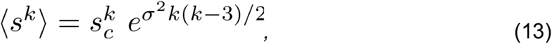

which, compared with the scaling prediction Eq. (3), is also consistent with *α* = 0, appropriate for the log-normal, and

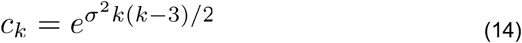

which is independent of 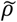 if *σ* is independent of 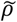, see Eq. (3).

This prediction is verified in **Figure 2B**, where the σ^2^ values are shown to be nearly uniform at different concentrations, with the exception of the data at the lowest concentration values, i.e. those furthest away from the critical concentration. These results are consistent with the scaling ansatz in the vicinity of 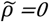.

Our analysis does not exclude the possibility that *σ* might depend on specific experimental conditions, even though the data that we analysed are suggestive of at most a weak dependence. We note that such dependence, even if present, does not invalidate the scaling, as long as the exponents do not depend on the experimental conditions.

### Estimation of the critical concentration of FUS using the scale invariance

The fact that the scaling ansatz is satisfied for different set of experiments opens a possibility to estimate the critical concentration. The scaling with φ=1 predicts 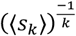 vs *ρ* to be a straight line with a slope depending on *k*. It is important to note that this is the consequence only of the scaling ansatz and not of the log-normal distribution. The line should intersect the *ρ*-axis at the critical concentration providing an estimate of *ρ*_*c*_.

We illustrate this process using the FUS data by plotting 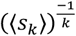 vs *ρ*. As expected, for different values of k we obtained a straight line fit of the points near *ρ*_*c*_ (**Figure 3**). Due to experimental uncertainties, the various lines, one for each value of *k*, lead to a slightly different estimate of *ρ*_*c*_. The average of the estimated *ρ*_*c*_ values is 5.0 ± 0.2 μM for FUS and 5.4 ± 0.4 μM for SNAP-tagged FUS (**Figure 3**). The different *ρ*_*c*_. values predicted based on the scaling ansatz using different k values enable the estimation of error of the predicted critical concentration (**Figure 3**). Both estimates are higher than the values of *ρ*_*c*_ originally reported (*ρ*_*c*_ = 2 μM of untagged and *ρ*_*c*_ = 3 μM of tagged FUS^27^) but consistent with them once taken into account the behaviour of the plateau of the absorbance of a spin-down assay used in that work. We also note that our estimates are compatible with other ones recently reported^33^.

**Figure 3:**
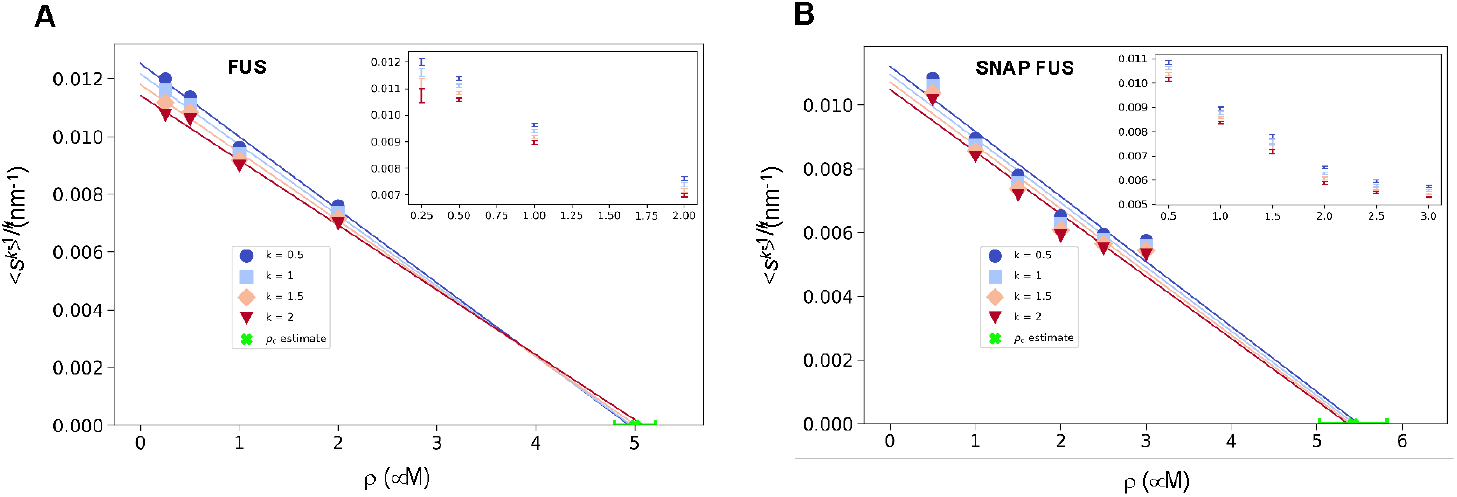
Estimation of the critical concentration of FUS using the scale invariance. **(A**,**B)** Critical concentration of FUS (5.0 ± 0.2 μM) (A) and SNAP-tagged FUS (5.4 ± 0.4 μM) (B). The scaling model predicts that the function of the moments plotted versus the concentration *ρ* becomes a straight line near the critical concentration *ρ*_*c*_ and intersects the *ρ*-axis at *ρ*_*c*_, independently of the value of *k*. Error weighted linear regressions are performed in both cases excluding the data point at the lowest concentration ρ = 0.125 μM. The resulting estimate of the critical concentration is shown in green along with the corresponding standard deviation, estimated from three independent measurements.

We then used both the values of *ρ*_*c*_ to probe the collapse of the size distribution functions (**Figure 4**). We observed that both in cases of untagged and tagged FUS, the SDFs collapsed with our estimated value of *ρ*_*c*_ (**Figure 4**).

**Figure 4.**
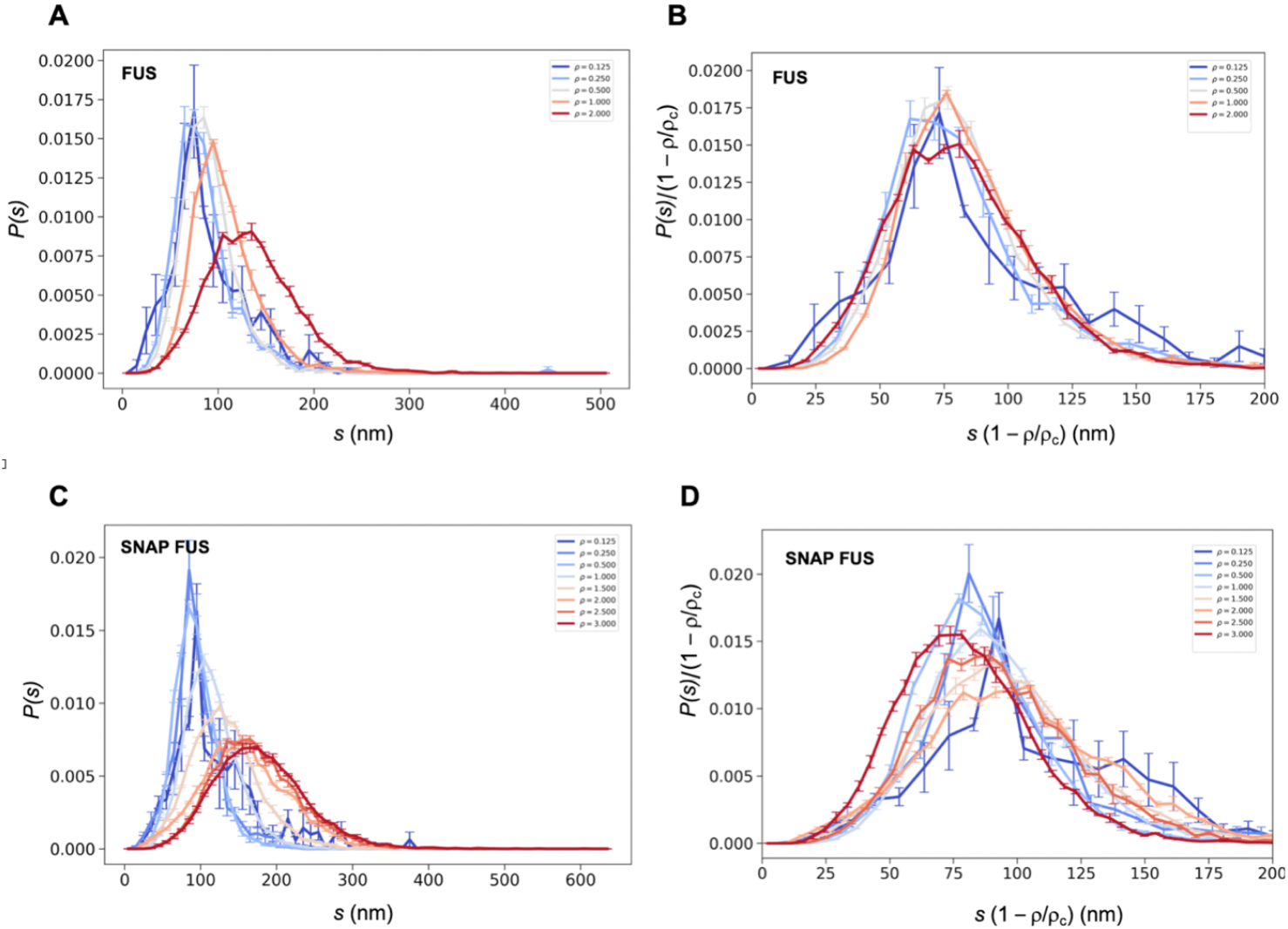
Collapse of the droplet size distributions of FUS as predicted by the scale invariance. If the scaling ansatz of Eq. (2) holds, the standard deviation σ of the log-normal distribution should not depend on the distance from the critical concentration, and a collapse should be achieved by rescaling the size with the distance 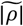 from the critical concentration. **(A**,**C)** Droplet size distributions derived from the experimental data of untagged FUS at 0.125, 0.25, 0.5, 1.0 and 2.0 μM concentrations (A) and SNAP-tagged FUS at 0.125, 0.25, 0.5, 1.0, 1.5, 2.0, 2.5 and 3.0 μM concentrations (C), and their standard error of the mean from three independent measurements (26). (**B**,**D)** Collapse of the droplet size distributions rescaled by the estimated critical concentration. The error bars show the standard error of the mean from the three independent measurements (26).

These results indicate that the scaling model can be used to estimate the critical concentration based on the distribution of droplet sizes.

### Estimation of the critical concentration of α-synuclein using the scale invariance

As α-synuclein droplets (called clusters) were recently reported below the critical concentration^34^, we aimed at characterising whether such droplets also follow scale invariant size distribution. Using A90C α-synuclein labelled with Alexa Fluor 647, we monitored droplet formation as previously described^35^. We measured droplet sizes at 5% PEG concentration 10 min after detection of liquid-like condensates using increasing concentrations of α-synuclein (20, 40, 50, 60, 75, 80 and 100 μM) (**Table S2**). Using k values of 0.25, 0.75, 1.25, 1.75, we determined the critical exponents in the scaling ansatz (Eqs. (4) and (6)), obtaining *φ* = 1.3 ± 0.2 (**Figure 5A**) and m=1-*α* = 0.9 ± 0.2 (**Figure 5B**). As control, we also determined the critical exponents using the scaling ansatz in a different way, using Eq. (19), obtaining *φ* = 1.1 ± 0.2 and m=1-*α* = 0.8 ± 0.2, corroborating the validity of the scale invariant model. We then estimated the critical concentration using the two methods obtaining *ρ*_*c*_ = 137 ± 10 μM, using Eq. (18) (**Figure 5C**) and *ρ*_*c*_ = 125 ± 7 μM, using Eq. (19) (**Figure 5D**). Images of the droplets at the different concentrations are shown in **Figure 6A**. We also observed that the droplet size distributions are stationary below the critical concentration, as expected (**Figure 6B,C**).

**Figure 5.**
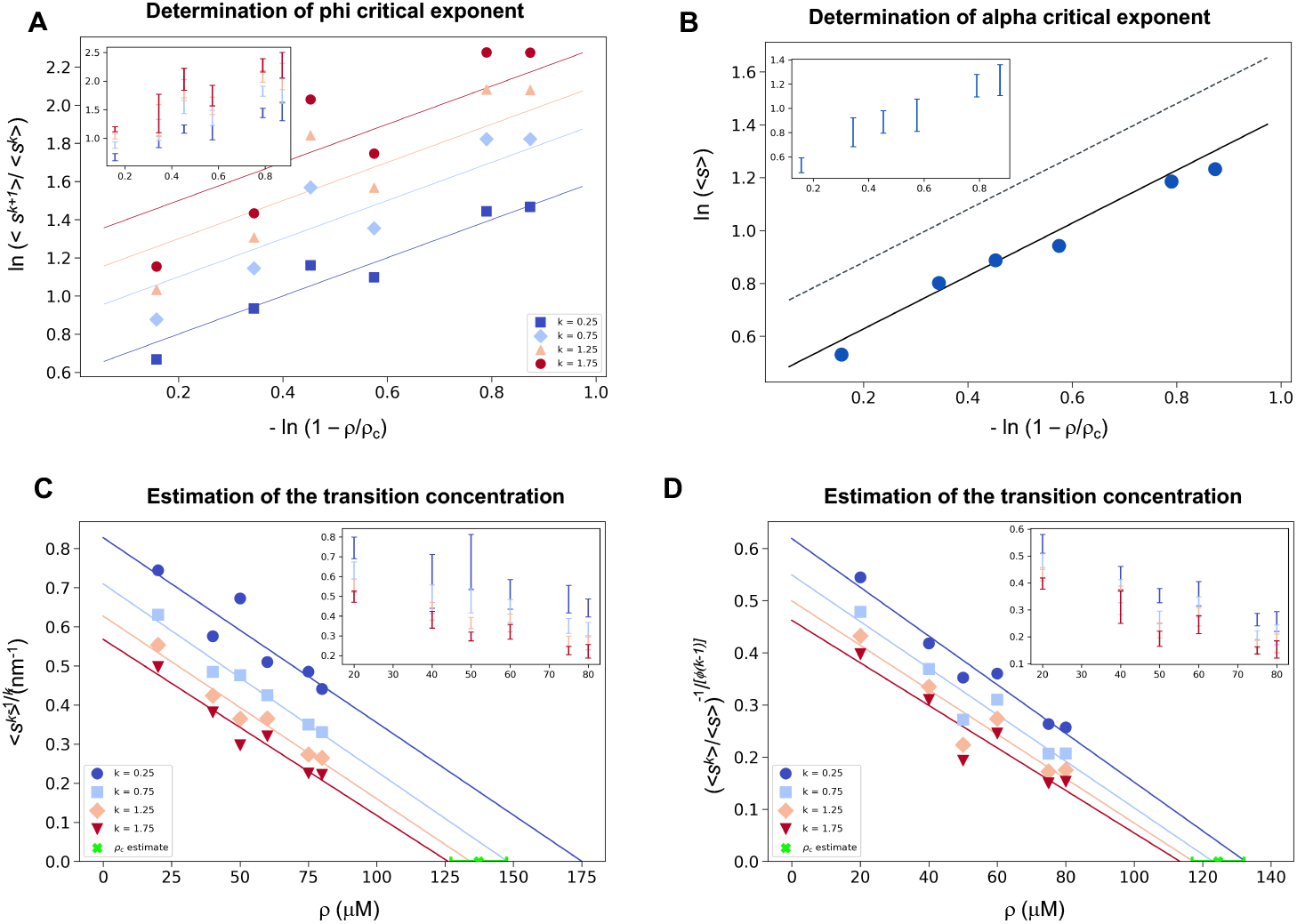
Estimation of the critical concentration of α-synuclein using the scale invariance. **(A)** Determination of the critical exponent *φ*. The ratios of the average moments of the droplet sizes (<s^k+1^>/<s^k^>, at k=0.25, 0.75, 1.25, 1.75, Eq. (4)) are represented at various distances 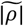 from the critical concentration. The exponent *φ* and its error for each value of k were determined as a mean and standard deviation of the three independent measurements (Eqs. (15) and (16)). Error bars are shown in inset for clarity. **(B)** Determination of the critical exponent m. The mean of the droplet size distributions is plotted at various distances 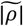 from the critical concentration. The value of the exponent *α* was determined by error weighted linear regression (Eq. 6), using φ=1, where the errors were standard deviations of the five independent measurements (Eq. (16)). Error bars, which were obtained as the standard deviation of the five independent measurements are shown in inset for clarity. Error-weighted linear regressions were performed. The fit corresponding to the scaling ansatz, compatible with *φ*=1 and *α*=0, is represented by a scattered gray line with a slope of 1. **(C**,**D)** Determination of the critical concentration for α-synuclein using the scaling ansatz in two different ways, either using Eq. (18) (C), resulting in *ρ*_*c*_ = 137 ± 10 μM, or Eq. 19 (D), resulting in *ρ*_*c*_ = 125 ± 7 μM, which are consistent within errors. The scaling model predicts that the function of the moments plotted versus the concentration *ρ* becomes a straight line near the critical concentration *ρ*_*c*_ and intersects the *ρ*-axis at *ρ*_*c*_, independently of the value of *k*. Error-weighted linear regressions were performed. The resulting estimates of the critical concentration are shown in green along with the corresponding standard deviation, estimated from four independent measurements. In panel D, φ was constrained to 1.0 using Eq. (19).

**Figure 6.**
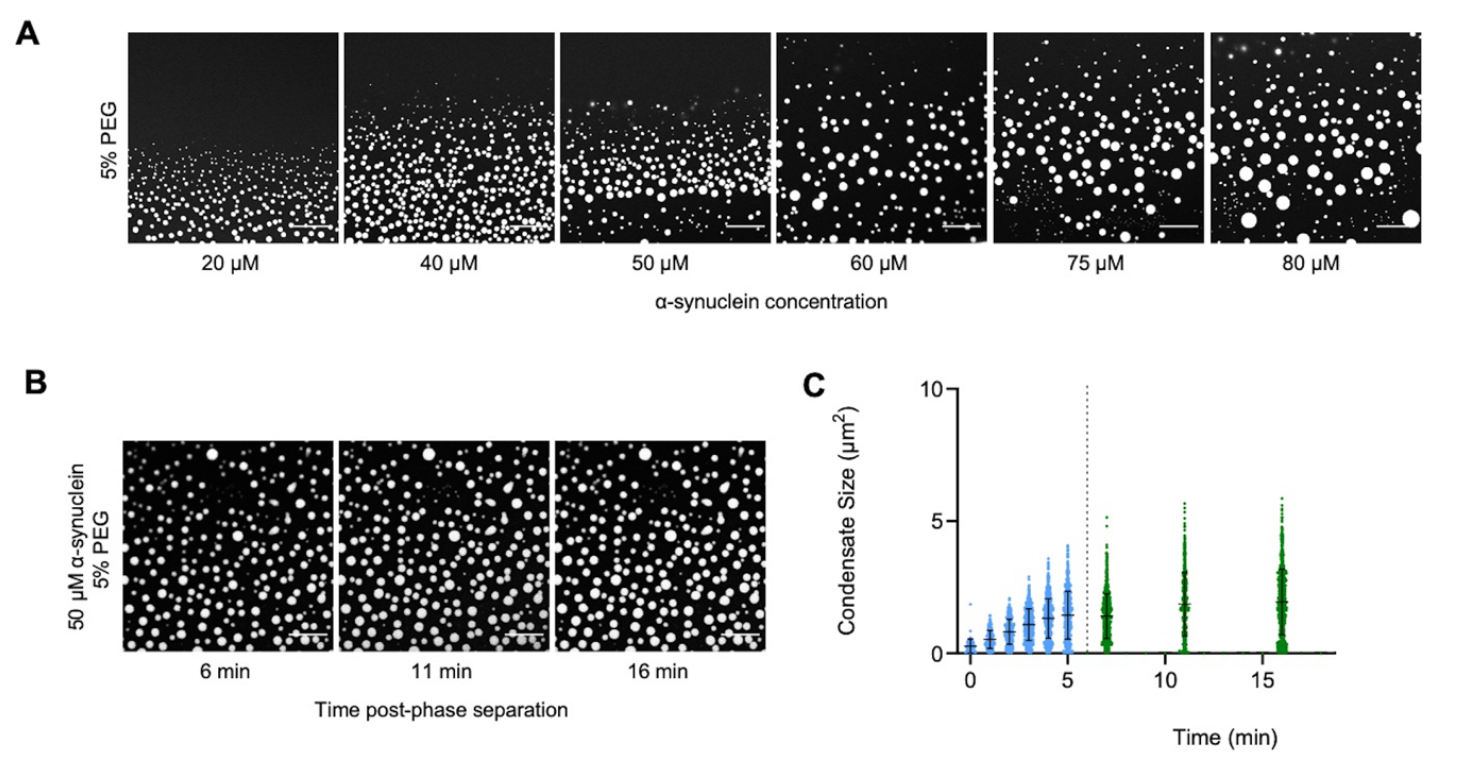
The droplet size distribution is stationary below the critical concentration. **(A)** Images of α-synuclein droplets at increasing concentrations of α-synuclein. **(B**,**C)** After an initial transient of 5 min, the droplet size distributions remain approximately stationary below the critical concentration, as shown for the case of 50 μM α-synuclein concentration.

## Discussion

Although growing experimental evidence indicates the presence of protein condensates both in vitro and in vivo^1-5,36^, their exact nature, and the mechanisms responsible for their formation are not fully understood. For example, it is unclear whether the droplets observed below the saturation concentration and those detected above result from the same process. Here we aimed at describing phase separation using a scaling ansatz.

With this aim, we analyzed a series of droplet size distributions of FUS and α-synuclein. We found that the droplet size distributions follow a scale-invariant log-normal behaviour. Scale invariance means that the description of the behaviour of a system remains the same regardless of the scale of observation. It has been seen for example in self-similar systems like fractals, which repeat patterns at different length scales^37^. Scale invariance is commonly observed in physics, biology, and economics, where it helps understand complex systems by identifying consistent patterns and fundamental properties^38^. The critical exponents of the scaling model are the same for different systems. We also note the generality of the log-normal model for nucleation and grain size growth ranging from crystal seeding^39,40^ to the mass size distributions in organism growth of various sea organisms^32^.

It is interesting to discuss the links of the scale invariance observed here with the theoretical models that have been proposed to date to explain the phenomenon of protein phase separation. A commonly adopted framework is the Flory-Huggins theory of phase separation^14-16^. In its simplest form, this theory describes a first-order phase separation in a system of homopolymers, which can also be adapted to polyampholites^17^. The Flory-Huggins theory has been extended to associative polymers, with the aim of modelling the sequence composition of proteins, by Flory and Stockmayer, who described the phase separation in terms of a third-order gelation process^18,19^. Semenov and Rubinstein modified the Flory-Stockmayer theory, reporting that the gelation process is not a real phase transition, as all the derivatives of the free energy are analytical at the gelation point^20^. It was also argued that another generalization of the Flory-Stockmayer theory^21^ could describe phase transitions in cytoskeletal networks^22^. In an older study, γ-crystallin was reported to undergo a second-order phase separation process^24^, consistently with observations in lysozyme solutions^25^.

More recently, it has been suggested that the protein phase separation process is coupled with percolation^23^. It is notable that percolation models exhibit a continuous transition marked by a diverging correlation length and a scale-invariant droplet size distribution^31^, resembling the general form as presented in Eqs. (1) and (2). However, the critical exponents for 3D percolation at the critical threshold are α=1.19 and φ=2.21, and in mean-field (d>6) scenarios, they are α=1.5 and φ=2, as per existing literature^31^. In contrast, our scaling analysis is consistent with α=0 and φ=1. Therefore, a phase separation coupled with percolation model^23^ could be consistent with the scaling behavior reported here if this coupling would induce a shift in the universality class from that of standard percolation^31^. This possibility should be further investigated to determine whether this type of models could predict the critical exponents α=0 and φ=1.

Scale invariance may appear to be in contrast with the Flory-Huggins theory, which characterizes droplet formation as a consequence of nucleation processes within a metastable state of a supersaturated system. In such cases, however, when the correlation length exceeds the cellular dimensions, the remnants of what should have been a first-order phase transition in an infinite system may effectively exhibit indistinguishable characteristics from a second-order phase transition. We also note that a power law distribution, but above the transition temperature, has recently been reported for nucleoli^41^.

Quite generally, the existence of scale invariance for the droplet size distribution, at least under the conditions investigated here, imposes stringent constraints on theoretical models that aim to elucidate protein phase separation. The importance of a scaling analysis lies in its ability to uncover the fundamental aspects of universal phenomena, transcending models confined solely to specific systems for which they were originally designed.

As a practical consequence of the scaling model, we found that the moments of the droplet size distribution can be used to obtain an estimate of the critical concentration for phase separation.

### Conclusions and Perspectives

It has been challenging to determine critical concentrations in the study of protein phase separation, particularly in cell systems. As a consequence, it remains still largely unclear when the observed protein droplets are formed at sub-saturation concentrations, and when they represent phase-separated condensates at super-saturation concentrations. To make progress towards understanding this problem, we have analyzed the probability distributions of the droplet sizes at different concentrations. The scale-invariant log-normal behaviour that we have reported offer a tool to determine the critical concentration. This approach should enable the positioning of a system with respect to the phase boundary when experimental data could be obtained about the length (1D), area (2D), or volume (3D) of the condensates. We anticipate that this method will find applications in the study of protein condensates under cellular conditions.

We also note that the scale invariance of the droplet size distribution that we have reported is characteristic of critical phenomena, which tend to be highly sensitive to environmental conditions. The formation and dimensions of protein droplets could therefore be rather tunable. This feature would appear to be favourable for the control of the formation of biomolecular condensates by the protein homeostasis system, and also suggest that protein phase separation could be efficiently modulated pharmacologically.

## Methods

### Expression and purification of α-synuclein

Human wild-type α-synuclein and A90C cysteine variants were purified from *Escherichia coli* BL21 (DE3)-gold (Agilent Technologies) expressing plasmid pT7-7 encoding for α-synuclein as previously described^35,42,43^. Following purification in 50 mM trisaminomethane-hydrochloride (Tris-HCL) at pH 7.4, α-synuclein was concentrated using Amicon Ultra-15 centrifugal filter units (Merck Millipore). The protein was subsequently labelled with a 1.5-fold molar excess of C5 maleimide-linked Alexa Fluor 647 (Invitrogen Life Technologies) overnight at 4 °C with constant mixing. The excess dye was removed using an Amicon Ultra-15 centrifugal filter unit and used immediately for phase separation experiments.

### Determination of the droplet distribution of α-synuclein

The experimental conditions were determined from a previously described phase boundary^35^. To induce liquid droplet/condensate formation, wild-type α-synuclein was mixed with an A90C variant labelled with Alexa 647 at a 100:1 molar ratio in 50 mM Tris-HCL and 5% polyethylene glycol 10,000 (PEG) (Thermo Fisher Scientific). The final mixture was pipetted onto a 35-mm glass-bottom dish (P35G-1.5-20-C; MatTek Life Sciences) and immediately imaged on Leica Stellaris Will inverted stage scanning confocal microscope using a 40×/1.3 HC PL Apo CS oil objective (Leica Microsystems) at room temperature. The excitation wavelength was 633 nm for all experiments. For liquid droplet size characterisation, images were captured 10 min post-liquid droplet formation. To analyse the possible time dependence of the liquid droplet size distribution, phase separation was induced, and 5 min post onset of phase separation, the glass-bottom dish containing the experiment was sealed and closed to maintain a stable and controlled environment. Subsequently, images were captured at each designated time point. All images were processed and analysed in ImageJ (NIH). Images were analysed by applying a threshold function in ImageJ that excluded the background of the image and identified the liquid droplets as having a circularity of 0.8-1.

### Determination of the droplet distributions of FUS

We used previously published data on FUS droplets^27^, where the fluorescence intensity of the FUS droplets was reported to be proportional to their diameters^27^. The abundance of untagged FUS droplets formed at 0.125, 0.25, 0.5 and 1.0, 2.0 μM concentrations were measured by nanoparticle tracking analysis for 51 different droplet sizes with three repetitions for each measure^27^. Similarly, the abundance of SNAP-tagged FUS droplets at 0.125, 0.25, 0.5, 1.0, 1.5, 2.0, 2.5 and 3.0 μM concentrations were measured for 64 different sizes with 3 repetitions for each measure^27^. For each data point, we determined the mean droplet abundance and the standard error of the mean from the three experimental replicates. The survival distribution function was computed as P_>x_ = N_>x_/N_tot_, where N_>x_ is the number of droplets above size x, and N_tot_ is the number of droplets. The values of x were chosen according to the original data^27^.

### Determination of the critical exponents for FUS

We computed the k-th moment of each droplet size, determined the average for each value of k and computed the ratios of the subsequent moments (Eq. (4)). We tested several k values from 0.25 to 3 in 0.25 increments, determined the moment ratios in each case. We selected four k values (k=0.5, k=1.0, k=1.5 and k=2.0) where the moment ratios at each concentration provided the best linear fit for both FUS and SNAP-tagged FUS. Then we generated a log-log plot of the moments and 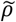, the distance from the critical concentration (**Figure 1**). For each value of k, the points were estimated by taking the average of the computation done on the 3 independent measurements. The errors of the data points are estimated as the standard error of the mean. At each k value, we performed a linear regression between the points at each concentration^27^ determining the slope, resulting in four φ_*k*_ values, with an associated error from the weighted linear regression. Given the independent observations φ_*k*_ with variance σ_*k*_, the value of the exponent was obtained as the weighted average

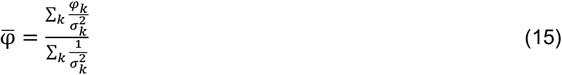

The error on the exponent φ was computed as

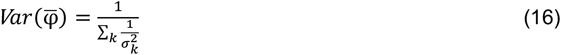

The exponent α was determined based on the log-log plot of the mean of the droplet size distribution at various distances from the critical concentration 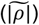. (**Figure 1**). We performed a linear regression weighted by the inverse of the variance of the data points corresponding, as previously, to the computed mean of the three independent experiments for each concentration (Eq. (6)) using φ =1. The error on the slope from the weighted linear regression is associated with the estimate of the exponent *m*. The exponent α is then derived using α = *1* − *m*.

At each concentration, from the distribution of ln(s) we determined s_0_ as <ln(s)> =ln(s_0_), and σ as the standard deviation of ln(s). At each concentration, we plotted the droplet size distribution function versus 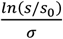 (Eq. (10)) using the s_0_ and σ values determined from the corresponding droplet size distribution at the given concentration. The log-normal behaviour is demonstrated by the resulting collapse (**Figure 2**). Furthermore, the collapsed curves overlapped with a theoretical log-normal curve computed with s_0_=1, σ = 1, which is the normal distribution in the rescaled log variables.

### Determination of the critical concentration for FUS

The *k*-th moment of the log-normal distribution (Eq. (13)) and the scaling ansatz with *α* = 0 (Eq. (3)) gives

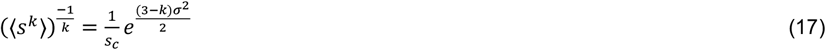

where 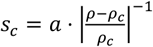, *a* is a proportionality constant independent of k, and σ is the standarddeviation of the logarithm of the droplet probability size distribution. Therefore

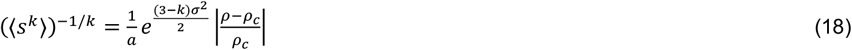

plotted versus the concentration intersects the x-axis at the value of the critical concentration.

### Determination of the critical exponents for α-synuclein

For α-synuclein, the critical exponents were determined in two different ways, the first using the same method as for FUS (Eq. (18)), and the second using

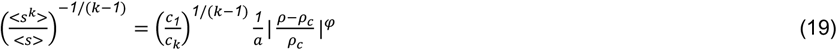

### Determination of the critical concentration of α-synuclein

As for the critical exponents, we determined the critical concentration of α-synuclein using the scaling ansatz in two different ways, either using Eq. (18) or using Eq. (19) with the constrain φ=1. We plotted the moments of the droplet size (Eq. (3)) versus the concentration, using a range of values of k to cover majority of the data. Analogously to what happens in Figure 3, the lines plotted in Figure 5D intercept the x axis on the same point, which corresponds to the critical concentration. We performed a linear regression weighted by the inverse variance, determining the intercept on the x-axis. For each value of k, we calculated the regression errors of both intercept and slopes. We estimated the critical concentration from each fit as the intercept calculated as 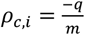, where *q* and *m* are the y-axis intercept and the slope retrieved by the fit, respectively, where the subscript *i* indicates each independent experiment. The estimated *ρ*_*c*_ was obtained as the mean of the different independent values *ρ*_*c,i*_. The error on the estimate of ρ_*c*_ is obtained as the standard error of the mean of the *ρ*_*c,i*_.

## Acknowledgements

We acknowledge insightful discussions with Drs. Serena Carra, Jonathan Vinet, Alex Buell and Soumik Ray. This work was supported by AIRC IG 26229 (M.F.).

